# The centrosomin CM2 domain is a multi-functional binding domain with distinct cell cycle roles

**DOI:** 10.1101/200204

**Authors:** Y. Rose Citron, Carey J. Fagerstrom, Bettina Keszthelyi, Bo Huang, Nasser M Rusan, Mark J S Kelly, David A. Agard

**Affiliations:** University of California, San Francisco; National Heart, Lung, and Blood Institute, National Institutes of Health; University of California, San Francisco & Chan Zuckerberg Biohub; University of California, San Francisco & Howard Hughes Medical Institute

## Abstract

The centrosome serves as the main microtubule-organizing center in metazoan cells, yet despite its functional importance, little is known mechanistically about the structure and organizational principles that dictate protein organization in the centrosome. In particular, the protein-protein interactions that allow for the massive structural transition between the tightly organized interphase centrosome and the highly expanded matrix-like arrangement of the mitotic centrosome have been largely uncharacterized. Among the proteins that undergo a major transition is the *Drosophila melanogaster* protein centrosomin that contains a conserved carboxyl terminus motif, CM2. Recent crystal structures have shown this motif to be dimeric and capable of forming an intramolecular interaction with a central region of centrosomin. Here we use a combination of in-cell microscopy and *in vitro* oligomer assessment to show that dimerization is not necessary for CM2 recruitment to the centrosome and that CM2 alone undergoes a significant cell cycle dependent rearrangement. We use NMR binding assays to confirm this intramolecular interaction and show that residues involved in solution interactions are consistent with the published crystal structure and identify L1137 as critical for binding. Additionally, we show for the first time an *in vitro* interaction of CM2 with the *Drosophila* pericentrin-like-protein that exploits the same set of residues as the intramolecular interaction. Furthermore, NMR experiments reveal a calcium sensitive interaction between CM2 and calmodulin. Although unexpected because of sequence divergence, this suggests that centrosomin-mediated assemblies, like the mammalian pericentrin, may be calcium regulated. From these results we suggest a model where during interphase CM2 interacts with pericentrin-like-protein to form a layer of centrosomin around the centriole wall and that at the onset of mitosis this population acts as a nucleation site of intramolecular centrosomin interactions that support the expansion into the metaphase matrix.

## Introduction

The centrosome is composed of hundreds of proteins that coordinate with one another to achieve a nucleation hub for microtubules that rearranges and duplicates as a function of the cell cycle. Until quite recently, little molecular detail has been available about the protein-protein interactions that regulate and organize this complex coordination. Several key proteins were identified decades ago via genetic interaction studies(1–4), including the *Drosophila melanogaster* proteins centrosomin (CNN), SPD-2, asterless, and pericentrin-like-protein (PLP), all of which are required for proper recruitment of the microtubule nucleation component γ-tubulin. Although once thought to be amorphous, super-resolution light microscopy has shown that the electron dense cloud that surrounds the centriole (peri-centriolar material or PCM) is organized into distinct protein domains (5–7). One set of proteins, including asterless and PLP, form a toroidal shape that remains close to the centriole wall throughout the cell cycle. A second set of proteins, including SPD-2 and CNN, also forms a ring around the interphase centriole but upon entrance into mitosis this expands outward from the centriole wall to form a much larger centrosome matrix containing the bulk of CNN and γ-tubulin. These distributions suggest the presence of discreet molecular interactions that define the interphase centrosome by tethering proteins, PLP, CNN, etc. at the centriole wall and that these proteins then act as a nucleation point for multivalent higher-order oligomerization of a subset of proteins at the onset of mitosis. However, this idea has not been fully confirmed and little detail is known on a molecular level about the interactions that would support interphase tethering or mitotic oligomerization. Recent work on *Caenorhabditis elegans* proteins (Spd2 and Spd5) has put forward a model of phase separation as an organizing principle for the mitotic centrosome (8). These contributions, along with yeast-two-hybrid data identifying a vast network of protein-protein interactions(9), begin to highlight the specific and elaborate protein organization within the PCM, although much of the underlying mechanistic detail is still lacking.

Of particular interest in this complex system is CNN as it takes on both a ring-like distribution in interphase as well as a matrix-like role in mitosis. CNN and its human homologue, CDK5RAP2, have consistently been identified as important for recruitment of the γ-tubulin ring complex (2,10,11) with the related self-assembling Spd5 playing an analogous role in *C*. *elegans*(12). Previous work has established that proper CNN recruitment is dependent on PLP(5), however, it was unclear whether this was through a direct interaction between the two proteins. More recent work using yeast-two-hybrid experiments has shown there is indeed a direct interaction between CNN’s carboxyl terminus and PLP(13) but no *in vitro* characterization has yet been performed. Meanwhile, crystallographic studies have revealed atomic details of a CNN intra-molecular interaction(14). Both these interactions have been attributed to a conserved motif at the carboxyl end of CNN, CM2. We sought to further probe CM2’s role in the centrosome in the hopes of understanding how it functions in these interactions and to shed light on the overall principles governing protein organization in the centrosome.

CNN contains two highly conserved motifs, the first of which, CM1, is near the amino terminus (Fig1A) and has been shown to interact with the γ-tubulin ring complex(11). The second motif, CM2, is positioned near the carboxyl terminus (Fig1A). Recent work by Feng *et al.* (14) revealed that *Drosophila* CM2 is dimeric, and identified key dimerization residues. Their crystal structures of CM2 show it is predominately helical and forms a complex with the middle portion of CNN (aa490-544) that contains a leucine zipper (CNNLZ). Here we confirm the dimerization *in vitro*, further define the requirements for dimerization, and analyze the implications of oligomerization for recruitment to the interphase and mitotic centrosomes in cells. By solution NMR we show that the CM2-CNNLZ complex forms in solution and that, *in vitro*, CM2 is multifunctional, with the ability to bind a region from the middle of pericentrin-like-protein (PLP) as well as calmodulin (CaM), suggesting that these interactions are mutually exclusive.

**Fig 1.**
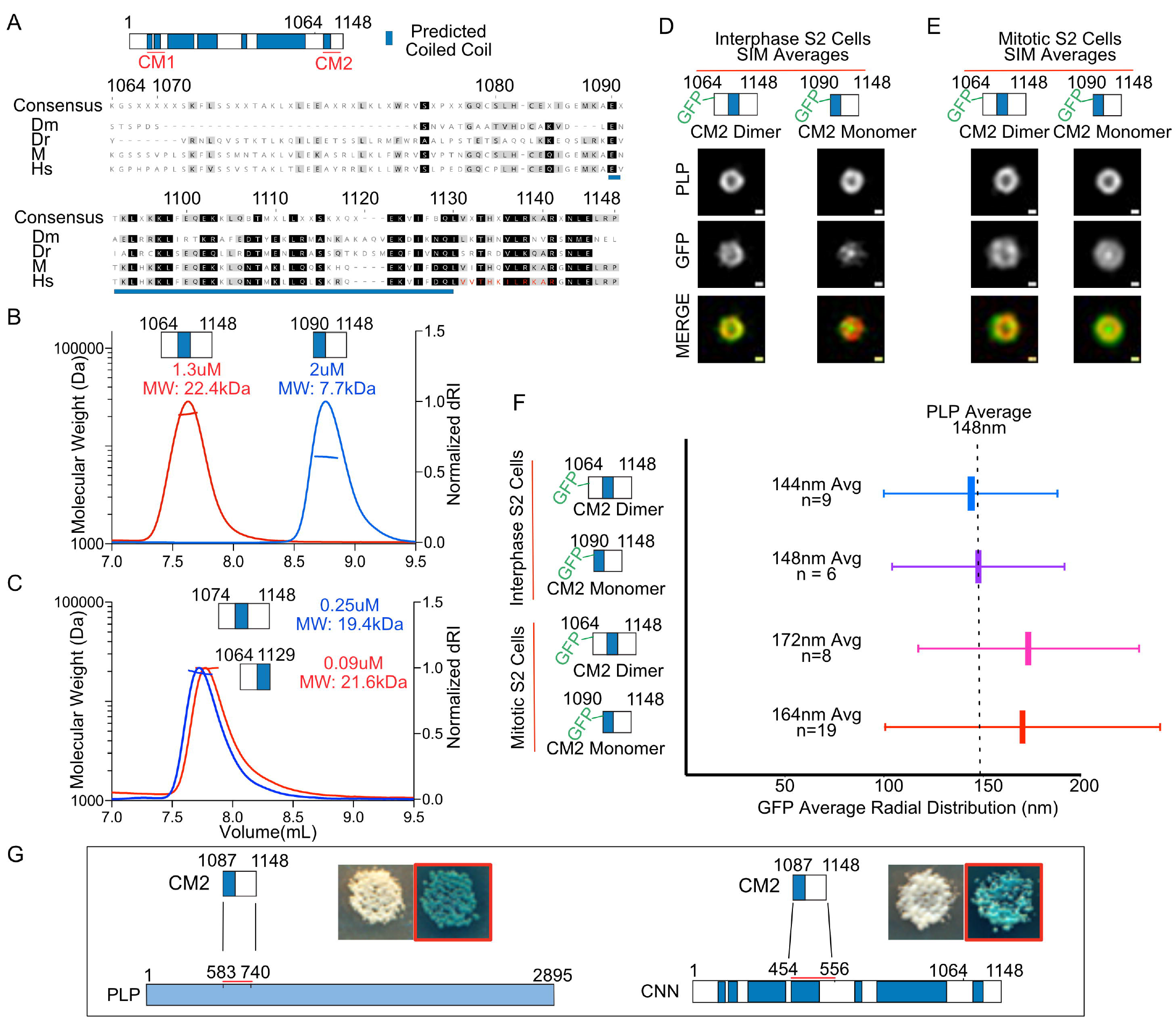
Conserved domain, CM2, of centrosomin characterized by SEC-MALS, SIM, and yeast-two-hybrid experiments. (A) Schematic of Centrosomin with coiled-coil prediction shown in blue. CM1 and CM2 are highlighted in red. Alignment of *D. Melanogaster’s* CM2 with the zebra fish, mouse and human (CDK5RAP2) homologues, with residue numbering according to *D. Melanogaster* residues. Higher conservation can be seen starting at residue E1090 in line with the start of the predicted coiled coil region. The predicted calmodulin binding site in human CDK5RAP2 is shown in red. (B) Molecular weights (left axis) measured with SEC-MALS are consistent with aa1064-1148 forming a dimer of CM2 (red) with a predicted monomer weight of 11kDa and a calculated weight of 22.4kDa. The construct aa1090-1148 exists as monomer (blue) with a predicted monomer weight of 7kDa and a calculated weight of 7.7kDa. (C) SEC-MALS of a more minimal dimer construct, aa1074-1148 (blue), that has a calculated molecular weight of 19.4kDa and predicted monomer weight of 9kDa. A construct with the final 18 residues truncated (Δ1130-1148) maintains a dimeric oligomer state (red) with a calculated molecular weight of 21.6kDa and predicted monomer weight of 8.8kDA. (D) Z-projections of the aligned and averaged PLP and GFP distributions for both the monomer and dimer GFP-CM2 fusion constructs in interphase S2 cells. Scale bar represents 100nm. (E) Z-projections of the monomer and dimer GFP-CM2 fusion constructs in mitotic S2 cells. Scale bar represents 100nm (F) Radial averages of the data represented in figures D and E with vertical bars indicating the average and horizontal bars representing the standard deviation of the Gaussian fit. (G) Schematic summaries of yeast-two-hybrid experiments showing the CM2 construct (aa1087-1148) interacting with a minimal domain of PLP (aa583-740) and the middle domain of CNN (aa454-556). Yeast plates of the interaction test are shown, with red boxes indicating plating on selection media.

## Results and discussion

To examine the ability of the c-terminus to form higher order oligomers and determine if CM2 alone can be recruited to the centrosome, two constructs were produced by recombinant expression in *E.coli* and purified. The first was designed to start at beginning of the predicted coiled-coil region in CNN that also coincides with highest sequence conservation (Fig1A,B) and extends to the C-terminus (aa1090-1148). The second construct (aa1064-1148) includes this same region but contains an additional 24 N-terminal residues that are not predicted to be coiled coil (Fig1A,B). This construct includes the residues identified by Feng *et al.* as being important for dimerization. Size exclusion chromatography with multi-angle light scattering (SEC-MALS) analysis of these two constructs reveals that while the shorter construct is monomeric, the addition of the 24 N-terminal residues results in dimer formation (Fig1B). Furthermore, a construct with intermediate boundaries (aa1074-1148) remains dimeric (Fig1C, blue) even at submicromolar concentrations. This construct was used as a dimeric construct for further *in vitro* studies while the longer dimer (aa1064-1148) was used for all further in cell assays. To test if the C-terminus contributes to dimerization, a construct with the final 18 C-terminal residues removed (CM2 Δ1130-1148) was assayed and was also found to be dimeric at submicromolar concentrations (Fig1C, red). These data, implicating residues 1074-1089 as being essential for dimerization, are consistent with Feng *et al.*’s observations and further suggests that the region of highest sequence conservation (aa1090-1148) is not important for dimerization but rather for interaction with partners.

To determine if both the monomeric and dimeric versions of CM2 are competent to be recruited to the centrosome throughout the cell cycle, GFP fusions of both constructs were made and transiently transfected into *Drosophila* Schneider 2 (S2) cells under the Sas6 promoter as previously described(15). Cells were co-stained with anti γ-tubulin antibodies and DAPI, and assessed for co-localization of GFP with γ-tubulin. In mitotic cells, both the monomeric and dimeric constructs are capable of being recruited to the centrosome with co-localization seen in 95% and 90% of cells, respectively.

Structured Illumination Microscopy (SIM) was used to assess GFP distributions in higher resolution at both interphase and mitosis. SIM images were aligned, averaged and fit with a Gaussian to obtain a radial average using previously described methods(5). At interphase both the monomeric and dimeric constructs have tight toroidal distributions similar to that of PLP (Fig1D,F), peaking at around 148nm, as would be expected for an interphase interaction between CM2 and PLP. On the other hand, during mitosis the monomeric and dimeric CM2 versions have both a larger radial average (164nm and 172nm, respectively) and a broader distribution (Fig1E,F) similar to full length CNN. These data suggests that CM2 alone is sufficient to integrate into the mitotic centrosome and, given the presence of endogenous CNN, is consistent with an intramolecular interaction between CM2 and the middle region of CNN. Though it had been known that full-length CNN underwent this change in distribution, our data newly indicate that CM2 itself is important for CNN’s recruitment both at interphase and at mitosis. We note that the averaged monomeric CM2 distributions at mitosis and interphase show a bright intensity at the center of the centrosome that does not appear in the dimer averages. While the origins of this central peak are unknown, the overall behavior indicates that a dimer interface is not necessary for recruitment although it may serve to increase the effective affinity. The striking difference in organization at these two cell cycle stages is suggestive of distinct molecular mechanisms, raising the possibility that two distinct binding partners maybe utilized to mediate recruitment at different times in the cell cycle.

Previous yeast-two-hybrid studies have been instrumental in identifying interaction partners critical for centrosome organization and for narrowing down the regions of interaction. These studies have shown that CM2 interacts with PLP aa583-1376 and that this interaction is not required for PLP recruitment to the centrosome but is essential for localizing PLP to centrosome flares within the PCM matrix (13). To narrow down the region of PLP interacting with CM2 and confirm that it is the CM2 region that is was responsible for both intra-and intermolecular CNN interaction, a refined yeast-two-hybrid screen was carried out. These experiments revealed that monomeric CNN was indeed capable of binding CM2 interaction partners and identified a minimal interaction region between monomeric CM2 and the central portion of CNN (residues 454-556, Fig1G). This is in agreement with Feng *et al.*’s observations of interaction between the same central fragment (termed CNNLZ) and dimeric CM2. Additionally, the yeast two-hybrid analysis narrowed down the PLP-CM2 interaction region to a short stretch of PLP (PLPMD, aa583-740, Fig1G). For a full list of constructs used in the yeast-two-hybrid experiments see supplemental figure 1.

To examine these interactions at higher resolution *in vitro*, the middle regions of CNN and PLP were fused to the GB1 solubility tag, purified and tested for CM2 interaction by observing changes in CM2 resonances in 2D [^15^N, ^1^H] NMR HSQC spectra. As a first step, the backbone resonances of monomeric CM2 (aa1090-1148) were assigned (BMRB: 27206) using standard triple resonance and residue-specific NMR experiments. Overlaying the HSQC spectra from monomeric and dimeric CM2 (aa1074-1148) reveals that the resonances of the final 40 C-terminal residues are unchanged between the monomer and dimer states (Fig2A,B), allowing the assignment of a significant number of residues in the dimer. From this result we infer that the structure of the C-terminal region of CM2 is unchanged upon dimerization, otherwise significant changes in chemical shift would have been observed. The region that does change in our spectra is quite consistent with the dimerization seen in the crystal structure (PDB 5MWE) of a complex between dimeric CM2 and a middle domain of CNN (Feng *et al*). Unfortunately, further interpretation of the dimer was hampered by significant variability in the dimer spectra that we attribute to multiple dimer species and/or multiple conformations. In support of this, both we (SFig2) and Feng *et al.* observed multiple dimer species by SEC-MALS at micromolar CM2 concentrations. Given that the monomer was sufficient in the two-hybrid studies, we used the monomer to explore solution interactions with binding partners.

**Fig2.**
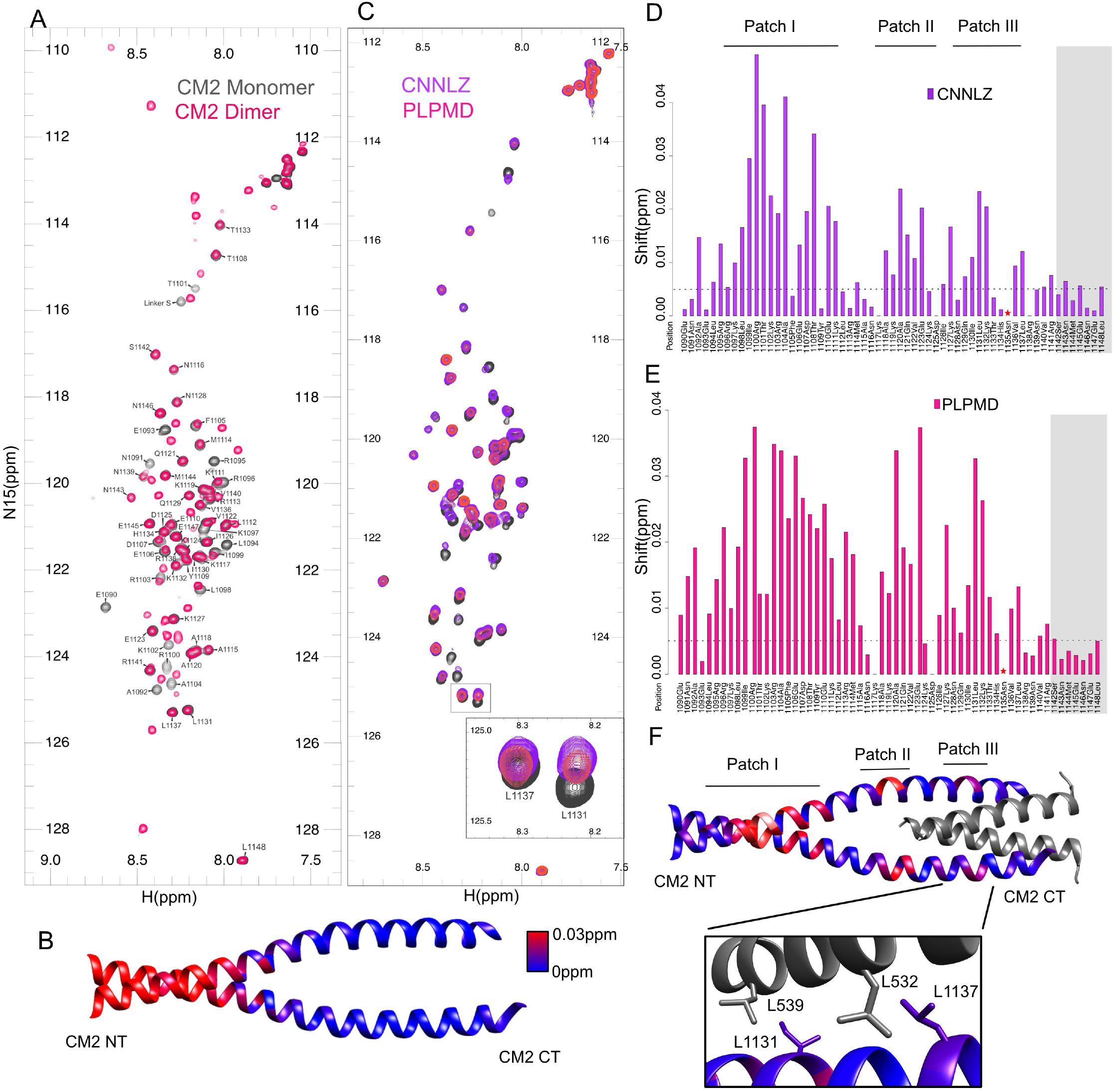
2D ^15^N-HSQC spectra of assigned monomeric CM2 shows interaction along three patches of residues. (A) Overlays of 2D ^15^N-HSQC spectra and assignments of CM2 monomer (gray) and dimer (pink) showing the peaks corresponding to residues F1105-L1148 have identical chemical shifts. All backbone monomer peaks are assigned. Though N1135 does appear in backbone tracing experiments, it does not appear in HSQC spectra. (B) Mapping of the combined chemical shift differences (blue to red) between the monomer and dimer HSQC spectra onto the crystal structure of CM2 (pdb:5MWE). Peaks that were not present in the dimer HSQC spectra were arbitrarily colored as a combined chemical shift difference of 0.1ppm for visualization purposes. (C) Overlay of monomer CM2 (gray) with either 200μM PLPMD (pink) or 150μM of CNNLZ (purple) illustrates chemical shift changes, particularly for L1131 and L1137 (inset). (D) Quantitation of the combined chemical shift changes of CNNLZ (purple) compared to monomeric CM2 alone reveals three regions with greatest shift changes. Dashed line indicates a combined chemical shift change of 0.0025ppm. The star indicates residue N1135 that is not visible in the HSQC spectra. Gray shading indicates the residues that are not modeled in the crystal structure (pdb:5MWE). (E) Quantitation of chemical shift changes of PLPMD (pink) compared to monomeric CM2. (F) Combined chemical shift changes of CM2 in the presence of CNNLZ mapped onto the crystal structure of CM2 with CNNLZ (pdb:5MWE). CM2 is colored blue to red in proportion to chemical shift changes. CNNLZ is shown in gray. Inset shows the interface between CM2 and CNNLZ and the proximity of I1130 to L539 and L1137 to L532. To assess the ability of monomeric CM2 to bind CNNLZ in solution we used the same CNNLZ fragment reported in the Feng *et al.* crystal structure (aa490-567, PDB:5MWE), while for the interaction assay with PLPMD we used the fragment defined by the yeast two-hybrid screen (aa583-740, Fig1E). 2D ^15^N HSQC spectra of monomeric ^15^N labeled CM2 (aa1090-1148) were collected in the absence or presence of 150μM CNNLZ (purple) or 200μM PLPMD (pink) (Fig2C). Addition of the GB1solubility tag alone showed negligible changes in the 2D ^15^N-HSQC spectra of CM2 (SFig3). In the presence of CNNLZ, multiple cross peaks are shifted and are concentrated into three patches when mapped onto the structure of CM2 in complex with CNNLZ (PDB 5MWE) (Fig2D/F). The first patch of residues (aa1092-1110) showing strong chemical shift perturbations is the very same region that changed upon dimerization. As previously noted, this also corresponds to the dimer contact observed in the crystal structure. Thus, binding to CNNLZ is sufficient to stabilize dimer formation even in the truncated CM2. The third patch is close to the C-terminus and includes several well-conserved hydrophobic residues (I1130/L1131/V1136/L1137). As shown by the crystal structure (Fig2F) these residues are in a region buried in the interface between CM2 and CNNLZ. In particular, we see shifts for the backbone ^1^H_N_, ^15^N resonances of I1130/L1131 and L1137, which are located in close proximity to CNNLZ’s L532 and L539 residues in the crystal structure (Fig2D, inset). The importance of these residues is also consistent with Feng *et al.*’s mutation analysis in cells showing that L532E and L539E mutants disrupt CM2’s recruitment to the pericentriolar material. Notably, the second patch is adjacent to the end of CNNLZ’s structured region, leading us to hypothesize that though not seen in the crystal structure, some portion of the remaining 23 residues of CNNLZ must be interacting with the CM2 patch II region. Interestingly, addition of PLPMD results in significant peak shifts for a remarkably similar set of CM2 residues. These residues map onto the same three patches (Fig2E), again indicating that PLPMD stabilizes CM2 dimerization. Although there is no structural information for PLPMD, it is predicted to contain a coiled-coil stretch (SFig4) according to the COILS prediction server (http://www.ch.embnet.org/software/COILS_form.html). Thus, like CNNLZ, PLPMD is presumably dimeric and contains a helical region that binds analogously to the same residues in CM2 as does CNNLZ. Similar experiments using dimeric CM2 in the presence of PLPMD and CNNLZ show significant reductions in peak intensities (SFig5), however, localizing those changes to particular residues is difficult for the reasons mentioned above. Together, these experiments indicate that two helical regions of CM2 not involved in dimerization form a binding surface for both intramolecular interactions (CNNLZ) and intermolecular interactions (PLPMD). Because the human homologue of CNN, CDK5RAP2, contains a predicted CaM binding site in its CM2 domain and immunoprecipitation experiments have confirmed this interaction (16), we asked if *Drosophila* CM2 might also bind CaM using the same NMR assay, despite the lack of a canonical CaM binding motif. In the presence of 20μM CaM and 2mM CaCl_2_ the CM2 HSQC spectra shows significant perturbation (Fig3A, pink). These shifts depend on CaM concentration, and even at concentrations as low as 5μM, peak shifts can be measured (Fig3B), suggesting that the affinity of CaM for CM2 is quite high and significantly greater than either *E. coli* produced CNNLZ or PLPMD. Mapping the chemical shifts onto the 5MWE crystal structure reveals shifts concentrated in the same patches as in the presence of CNNLZ and PLPMD, as well as additional shifts along the length of the helices. As the shifted regions are also consistent with dimer interactions, we speculate that CaM is even more potent at dimerizing monomeric CM2 than either CNNLZ or PLPMD. Our NMR chemical shift perturbations indicate CaM binds the hydrophobic residues at the CM2 C-terminus. This is also consistent with the known requirement for hydrophobic residues in calcium dependent CaM binding, though the patterning of hydrophobic residues in CM2 is not ideal (17) To assess the effect of calcium on binding, experiments were performed with 2mM EGTA added in place of 2mM CaCl_2_ (Fig3A, purple). Under these conditions the chemical shift perturbations are significantly reduced but are not completely absent. Based on comparison of chemical shifts under EGTA conditions and titrations of CaM in the presence of calcium, we estimate that the apparent affinity in the absence of calcium is reduced by 50-70%. As with PLPMD and CNNLZ binding, ^15^N labeled dimeric CM2 shows very significant reductions in peak intensity upon CaM addition for a number of residues (SFig6), which are likely due to cross peak broadening. Though these experiments support a calcium dependent CaM interaction, gaining a better understanding of the mechanism and function of this interaction will require further structural and cellular studies.

**Fig. 3.**
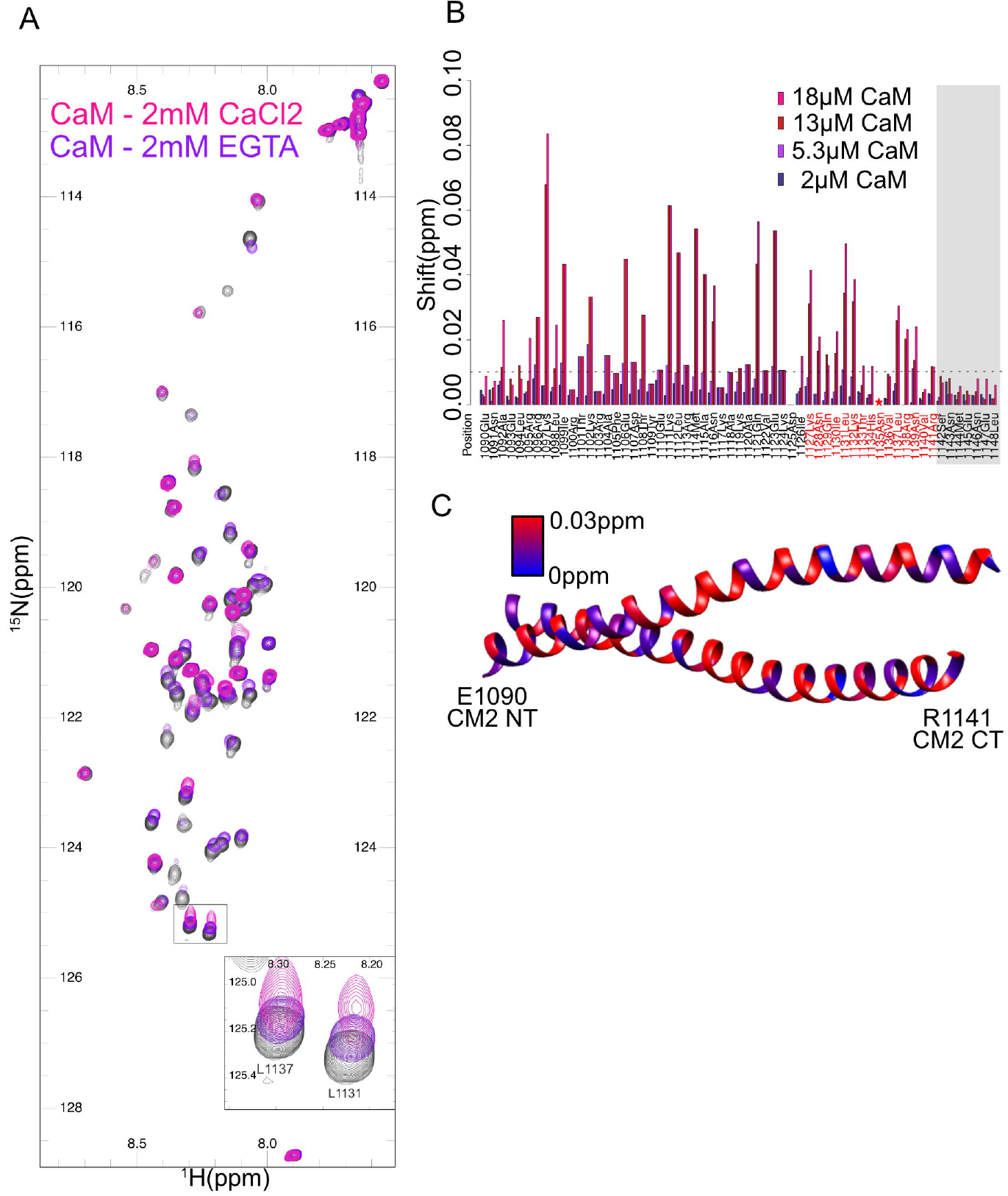
2D ^15^N-HSQC spectra of monomeric CM2 show a calcium sensitive interaction of calmodulin. (A)2D ^15^N-HSQC spectra of monomeric CM2 alone (gray) overlaid with the spectra after the addition of 18μM CaM with 2mM CaCl_2_ (pink) shows significant shifts, while 18μM CaM in the presence of 2mM EGTA (purple) shows more minimal shifts. Inset: Shifts on leucine 1131 and leucine 1137 are apparent. (B) Quantitation of shifts from a CaM titration (2mM CaCl_2_) where several shifts are greater than 0.01ppm (dashed line) on addition of 5.3μM CaM. Residues that align with the predicted CaM binding site in mammalian CDK5RAP2 are shown in red. The star indicated residue N1135 that is not visible in the HSQC spectra while the gray shading indicates residues that are not modeled in the 5MWE crystal structure. (C) Mapping of chemical shift changes (blue-red) onto the 5MWE crystal structure (CNNLZ is shown in gray) reveals shifts along the length of the helices including the region where dimer contacts are made. To interrogate in greater detail the functional contributions of specific regions and residues implicated by the HSQC binding data, truncations and mutations were designed in GFP fusions of dimeric CM2 and assessed in S2 cells. Firstly, to determine if the most carboxyl terminal residues were required for recruitment, a truncation at residue 1130 was made in the GFP-dimeric CM2 construct (Fig4A, CM2 Δ1130-1148) and localization to the centrosome was completely abolished (Fig4B). Maximum GFP pixel intensities in regions identified as the centrosome by γ-tubulin staining show an 85% reduction in maximum intensities when compared to the wildtype CM2 construct (Fig4B). By contrast, removal of the final 7 residues (CM2 Δ1141-1148) had no effect on centrosome recruitment, despite their high level of conservation. This is consistent with our HSQC experiments showing minimal changes in this region upon CNNLZ binding (Fig2C,E & 3B). To investigate the role of the intermediate residues, alanine scanning was performed in blocks of four residues across the relevant region (aa1130-1141). While mutating the span RNVR (Fig4A) to alanines did not reduce the ability of CM2 to localize at the centrosome, mutating either the ILKT block or the HNVL block significantly reduced recruitment by 29% and 55%, respectively (Fig4B). These data are in agreement with our HSQC data that show significant shifts in leucines 1131 and 1137 that are contained within these spans. Furthermore, a single point mutant replacing leucine 1137 with an alanine nearly abolished recruitment, reducing the maximum pixel intensity by 80% (Fig4B,C). Importantly, a more conservative mutation of L1137 to a valine shows a partial recovery of recruitment, suggesting a critical hydrophobic interaction is mediated by this residue (Fig4B).

**Fig. 4.**
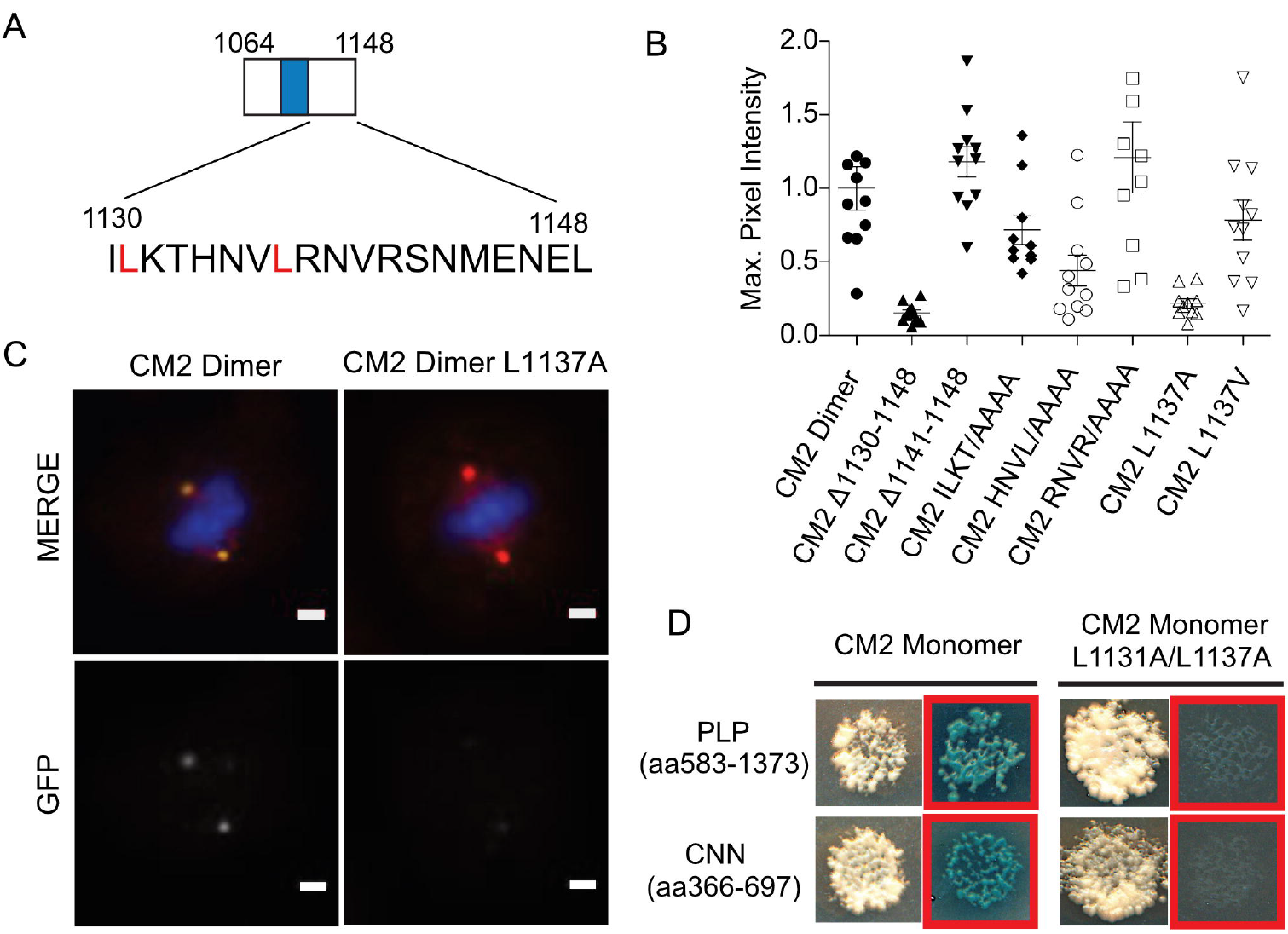
Leucines 1131 and 1137 implicated as important for CM2 interaction with CNNLZ and PLP. (A) Schematic of dimeric CM2 with residues 1130-1148 highlighted. (B) Quantification of the maximum GFP pixel intensity at the centrosome normalized to dimeric CM2 (aa1064-1148). From the left, GFP-CM2 dimer fusion serves as the control to which all other GFP constructs are compared. Truncating off the final 18 residues, CM2 Δ1130-1148, shows an abolishment of recruitment to the centrosome while truncating the final 7 residues, CM2 Δ1141-1148, has no apparent effect on recruitment. Alanine mutant constructs of ILKT to AAAA and HNVL to AAAA have 28% and 56% reduction in GFP intensity respectively, while RNVR to AAAA appears unperturbed. Single point mutations of L1137 to alanine and valine have 78% and 27% reductions, respectively. (C) Representative wide field microscopy images of wildtype CM2 dimer (left) with identical scaling and contrast of CM2 dimer L1137A point (right) mutation images. The top row shows merging of DAPI (blue), gamma tubulin (red) and GFP (green) while the bottom is only the GFP channel. 1um scale bar. (D) Yeast-two-hybrid plates comparing monomeric CM2 prey – PLPMD bait colonies and monomeric CM2 prey – CNNMD bait colonies (right) with identical experiments using CM2 double mutant L1131A/L1137A (left). Red boxes indicate plating on selection media. Consistent with these observations, yeast-two-hybrid data with a double L1131A/L1137A mutant show abolishment of the interaction with CNNLZ, but also extended our studies by showing a loss of interaction with PLPMD (Fig4D). Notably, the yeast-two-hybrid experiments were done with fragments of CNN middle domain and PLPMD significantly larger than the determined minimal domains (Fig1F) indicating that secondary regions are not contributing to the interaction by contacting CM2 outside patch I. Though Feng *et al.* argue that I1126 and T1133 are also required for interaction with CNNLZ, we see only a minimal perturbation of CM2 recruitment when ILKT is replaced with alanines. This suggests that Feng et al.’s mutation to glutamic acid is strongly disruptive of the hydrophobic pocket required for interaction but that these residues are not actively involved in contacting CNNLZ. Rather, we show that mutation of leucine 1137 alone can abolish recruitment in cells, an observation that is fully consistent with our NMR data showing chemical shift changes for this residue when CM2 binds all three of its interaction partners. Taken together, our yeast-two-hybrid, cellular microscopy, SEC-MALS experiments and NMR data support a model where CM2 plays distinct roles in interphase and mitosis through interactions with several binding partners and that these are ultimately responsible for the different phases of PCM establishment and expansion. We suggest that during interphase CNN is tethered near the centriole wall via the interaction of CM2 with PLPMD and that upon transition into mitosis the CNNLZ domain of this tethered protein serves as a nucleation point for the formation of a multivalent higher-order assemblies mediated primarily by CM2- CNNLZ interactions, likely in a phosphorylation-dependent manner. Our data suggest that both these interactions exploit the same binding mode with CM2 in order to interact through the conserved residue L1137. Though monomeric CM2 is sufficient to bind both CNNLZ and PLPMD, given the importance of this single residue for both interactions, it is likely that dimeric CM2 is favored in cells by providing increased affinity and refining localization. Furthermore, these data suggest that PLPMD is likely to be a helical dimer that presents itself to CM2 similarly to CNNLZ. Though we show convincingly that CM2 is also capable of interacting with CaM in a calcium dependent manner, uncovering the functional role of this interaction will require further investigation. Nevertheless, given the high affinity of CaM for CM2 and that HSQC spectra implicate the same hydrophobic residues for all three interactions tested here, CaM may function as a regulatory mechanism that modulates PCM assembly by competing with PLPMD or CNNLZ for binding to CM2.

## Materials and methods

### Protein expression and purification

CNN cDNA (LD19135) was obtained from the DGRC and sub-cloned into the 6xHIS containing pET28a expression vector modified to replace the TEV cleavage site with a PreCission protease cleavage site. For unlabeled protein, *E-coli* was grown in terrific broth at 37°C to OD_600_ - 0.6-0.8 then induced with 1mM IPTG, cooled to 18°C and grown for 16-18hours before harvesting. Samples labeled with ^15^N or ^13^C were grown in 3g/L KH_2_PO_4_, 6g/L Na_2_HPO4, 0.5g/L NaCl, 0.011g/L CaCl_2_, 0.18g/L MgSO4, 0.001g/L thiamine, 0.001g/L biotin with the addition of either unenriched or labeled ammonium salt and/or glucose as a carbon source at 1.25g/L (^15^NH)SO and 2g/L ^13^C glucose. Cells in labeling media were grown to OD_600_ - 0.5 then induced with 1mM IPTG, cooled to 18°C and grown for 16-18hours before harvesting.

For all proteins, cells were harvested by centrifugation then pellets were either frozen for later use or lysed immediately with a Emulsiflex C3 in 50mM HEPES pH 7.0, 300mM NaCl, 10mM imidazol, and 6mM beta-mercaptoethanol. Lysates were clarified by ultracentrifugation at 35,000xg and the supernatant was incubated with Qiagen NiNTA Superflow resin with gentle agitation for 1.5-2 hours. The resin was washed in batch using 50ml lysis buffer and eluted with a 300mM imidazol buffer. The 6xHIS tagged was cleaved overnight with 3C protease (except in the case of CM2 Δ1130-1148). The pooled elution was then further purified by cation exchange chromatography. Additionally, non-labeled proteins were further purified by on size exclusion columns.

## SEC-MALS

Multi-angle light scattering analysis was performed with a KW-802.5 size exclusion column on an Ettan liquid chromatography system (GE Healthcare Life Sciences) with inline DAWN HELEOS MALS and Optilab rEX differential refractive index detector (Wyatt Technology). All samples were in 50mM HEPES pH 7.0, 200mM NaCl, and 1mM DTT running buffer. Data were analyzed using ASTRA VI system software (Wyatt Technology).

### S2 cell microscopy sample preparation

*Drosophila* Shnieder 2 (S2 cells) from ATCC were transfected with PMT backbone plasmids using Amaxa Cell Line Nucleofector Kit V (Lonza). After 34-38 hours dimer-CM2 and dimer-CM2-L52A underwent fluorescence flow cytometry and showed equivalent GFP expression but were sorted nonetheless and plated. All other cells were plated after 36 hours on convanavalin-A (sigma) coated glass bottom dishes. Samples for SIM were plated on 35mm glass bottom Delta T dishes (Bioptechs). After 3 hours cells were washed with PBS and fixed in cold methanol for 20min. Cells were then rehydrated in PBS and blocked in PBS with 0.02% tween and 3% BSA. Samples for wide field microscopy were stained 1:500 with rabbit anti-GFP antibodies (Invitrogen) and 1:75 mouse anti γ-tubulin (Sigma GTU88). SIM samples were stained with 1:500 chicken anti-GFP antibodies (Invitrogen) and 1:100 anti-PLP (86-PPH-A against aa683-974, a kind gift from Jordan Raff). Fluorescent secondary antibodies Alexa 488 or Alexa 555 were then applied to the sample at 1:500 as well as 300nM DAPI. All samples were mounted in an v/v 90% glycerol, v/v 10%Tris pH 8.0 and w/v 0.5% propyl-gallate media.

### Yeast-two-hybrid analysis

Cnn 366-697, Cnn 454-556, Cnn 1087-1148, PLP 583-1373 and PLP 583-740 pieces were amplified from cDNA clones by PCR using Phusion (Thermo Fisher Scientific, Waltham, MA) and with the primers

Cnn 366_F CACCATGTCCTCCAGCGGCCGTTCCATGAGTGAC,
Cnn 697_R AGGCGCTGCCAAACTGTTGGAGGTTTC,
Cnn 454_F CACCATGGAAGCAGATCTGCAGCAATCCTTCACGGAG,
Cnn 556_R TCGATCAGCGGCCAACACGCC,
Cnn 1087_F CACCATGGTAGATCTTGAAAACGCC,
Cnn 1148_R TAACTCATTCTCCATGTTTGAGCGAAC,
PLP 583_F for PLP 583-1373 CACCATGTCCCTCTCCTTGGATGAGTC,
PLP 1373_R TGGAGGTAGGGAGGAATGTGTTTTTCCC,
F- BamHI NdeI 583 for PLP 583-740
CACCATGGGATCCCATATGATGCCTTCCCTCTCCTTGGATGAG
PLP 740_R GCGAGTCCTGCGGCCGCTTAACTCGAGCGTTTCACTGCATCG.

PCR products were then introduced into Gateway Entry vectors using the pENTr/D-TOPO Kit (Thermo Fisher Scientific). All Y2H experiments were then conducted as described previously in Galletta et al 2016 (ref: Galletta BJ, Fagerstrom CJ, Schoborg TA, McLamarrah TA, Ryniawec JM, Buster DW, Slep KC, Rogers GC, Rusan NM. A centrosome interactome provides insight into organelle assembly and reveals a non-duplication role for Plk4. Nat Commun. 2016 Aug 25; 7: 12476.). In brief, fragments of Cnn and PLP were introduced into pDEST-pGADT7 and pDEST-pGBKT7 using Gateway technology (Thermo Fisher Scientific), transformed into Y187 or Y2HGold yeast strains (Takara Bio USA, Mountain View, CA), and grown in −*leu* or −*trp* media to select for plasmids. After mating of the two strains, yeast were grown on −*leu*, −*trp* (DDO) plates, then replica plated onto plates of increasing stringency: DDO; −*ade*, −*leu*, −*trp*, −*ura* (QDO); −*leu*, −*trp* plates supplemented with Aureobasidin A (Takara Bio USA) and X-α-Gal (Gold Biotechnology, St. Louis, MO) (DDOXA); and −*ade*, −*leu*, −*trp*, −*ura* plates supplemented with Aureobasidin A and X-α-Gal (QDOXA). Interactions were scored based on growth and the development of blue color as appropriate. All plasmids were tested for the ability to drive reporter activity in the presence of an empty vector (autoactivation). Plasmids that conferred autoactivity were omitted from further analysis.

### Microscopy acquisition and analysis

SIM data was acquired on a DeltaVision OMX SR microscope (GE Healthcare) using a 60×/1.42 NA oil immersion PSF objective and three sCMOS cameras. Immersion oil with a refractive index of 1.514 was used to match with the refractive index of the sample. Z stacks of 9μm thickness were collected with 0.125μm step-sizes using 5 phases and 3 angles per image. Reconstructions were preformed using SoftWorx 6.5.2 (GE Healthcare) with a 0.003 Wiener filter for the Alexa Fluor 488 channel and 0.002 Wiener filter for Alexa Fluor 555 as suggested by analysis using the ImageJ plugin SIMcheck (18). Channel specific optical transfer functions were used for reconstructions. Alignment, averaging and calculation of the radial distribution were performed using custom python scripts previously described(5). Monomeric CM2 data was fit with an additional central Guassian to distinguish the central intensity from the ring distribution.

Widefield microscopy was acquired on a Ziess Axiovert 200 and GFP intensity was quantified using Matlab. Experiments that were not done in single batch were normalized using GFP quantification of GFP-Dimer CM2.

### Nuclear magnetic resonance

Double labeled (^13^C/^15^N) protein was expressed as described above with ^15^N ammonium chloride and ^13^C_6_ glucose and purified. The following 3D triple resonance experiments for backbone assignments were recorded on a 500MHz Bruker AVANCE DRX500 spectrometer equipped with a Z-gradient QCI inverse cryoprobe (^15^N/^13^C/^31^P,^1^H): CBCANH (pulse program: cbcanhgpwg3d), CBCAcoNH(pulse program: cbacaconhgpwg3d), and hNcaNHn (pulse program: hncannhgp3d). NMRPipe was used to process 3D spectra(19). Proton chemical shifts were referenced to an external DSS standard and ^15^N and ^13^C shifts referenced indirectly from this value. Double labeled protein was also used to perform the following multiplicity selective in-phase coherence transfer (MUSIC) experiments on a 800 MHz Bruker AVANCEI spectrometer equipped with a Z-gradient TXI cryoprobe: music_qn_3d, music_de_3d, music_ile_3d, music_cm_3d, music_tavi_3d, music_kr_3d(20). All experiments were performed at 300K in 10mM Trizma pH 7.2, 150mM NaCl, 0.01%(w/v) NaN_3_ and 5% (v/v) D_2_O.

All 2D [^15^N, ^1^H]-HSQC (pulse program: fhsqcf3gpph) binding experiments were carried out with 100μM of ^15^N labeled protein on an 800 MHz Bruker AVANCEI spectrometer. HSQC experiments were performed at 300K with all proteins dialyzed overnight into 20mM PIPES pH 6.8, 140mM NaCl, 0.01% (w/v) NaN_3_, 5% (v/v) D_2_O and unless otherwise noted contained 2mM CaCl_2_.

Resonance assignments and analysis was done in CCPNMR(21). Peak intensities and shifts were analyzed and plotted in R. Combined chemical shift were calculated using the formula Δδ(^1^H) + Δδ(^15^N)/5 as described by Hajduk et *al*.(22).

## Acknowledgments

We thank members of the Huang and Agard lab for helpful discussions. We thank Wei Qiang and Veronica Pessino in the Huang lab for help with cell sorting at various stages of the project. Furthermore, we thank Jeffery G. Pelton for help and access to the Central California 900 NMR facility as well as helpful advice from James Holton at early stages of the project. We thank Jordan Raff for the anti-PLP antibody.

## Supporting information

**SFig1. Schematic of all yeast-two-hybrid experiments.**

**SFig2. SEC-MALS of CM2 aa1074-1148 indicates multiple dimer species.**

CM2 construct used for NMR appears dimeric by SEC-MALS analysis. At 1μM (blue) this construct appears a predominately, as a single dimeric peak with a calculated molecular weight of 19.2kDa while the predicted monomer weight is 9kDa. At 9μM (red) this same construct appears as three or more not fully resolved peaks, however, the molecular weight across all these peaks appears to remain dimeric.

**SFig3. GB1 control 2D ^15^N-HSQC experiment.**

(A) HSQC spectra from a control experiment of monomeric CM2 (gray) and monomeric CM2 with 150μM GB1 (cyan) overlaid. All the peaks appear unperturbed in the presence of GB1. B) Quantitation of peak shifts on addition of GB1 with the same y-axis scaling as for the GB1-CNNLZ and GB1-PLPMD data in Fig2D,E.

**SFig4. Coiled-coil prediction of PLPMD.**

Coiled-coil prediction of PLPMD from the COILS server shows high probability of a coiled coil secondary structure in the carboxyl region of PLPMD.

**SFig5. 2D ^15^N-HSQC experiments of dimeric CM2 in the presence of CNNLZ and PLPMD.**

HSQC spectra (gray) of dimeric CM2 with low intensity peaks at lower (<8ppm) proton resonances indicative of multiple species. Overlaid are dimeric CM2 with addition of 75μM CNNLZ (purple) and 100μM PLPMD (pink) that exhibit reduced intensities on a significant number of peaks. All spectra are contoured and scaled identically.

**SFig6: 2D ^15^N-HSQC experiments of dimeric CM2 in the presence of CaM**

Overlay of the HSQC spectra of dimeric CM2 (gray) and dimeric CM2 with the addition of 18μM CaM (pink) in the presence of 2mM CaCl_2_. Both spectra are contoured and scaled identically. The intensities of several peaks are reduced dramatically with the addition of CaM and new peaks also appear in the presence of CaM.

